# Does pericentral mu-rhythm “power” corticomotor excitability? – a matter of EEG perspective

**DOI:** 10.1101/2020.09.11.292789

**Authors:** Anke Ninija Karabanov, Kristoffer Hougaard Madsen, Lærke Gebser Krohne, Hartwig Roman Siebner

## Abstract

**Background:** Electroencephalography (EEG) and single-pulse transcranial magnetic stimulation (spTMS) of the primary motor hand area (M1-HAND) have been combined to explore whether the instantaneous expression of pericentral mu-rhythm drives fluctuations in corticomotor excitability, but this line of research has yielded diverging results.

**Objectives:** To re-assess the relationship between the mu-rhythm power expressed in left pericentral cortex and the amplitude of motor potentials (MEP) evoked with spTMS in left M1-HAND.

**Methods:** 15 non-preselected healthy young participants received spTMS to the motor hot spot of left M1-HAND. Regional expression of mu-rhythm was estimated online based on a radial source at motor hotspot and informed the timing of spTMS which was applied either during epochs belonging to the highest or lowest quartile of regionally expressed mu-power. Using MEP amplitude as dependent variable, we computed a linear mixed-effects model, which included mu-power and mu-phase at the time of stimulation and the inter-stimulus interval (ISI) as fixed effects and subject as a random effect. Mu-phase was estimated by post-hoc sorting of trials into four discrete phase bins. We performed a follow-up analysis on the same EEG-triggered MEP data set in which we isolated mu-power at the sensor level using a Laplacian montage centered on the electrode above the M1-HAND.

**Results:** Pericentral mu-power traced as radial source at motor hot spot did not significantly modulate the MEP, but mu-power determined by the surface Laplacian did, showing a positive relation between mu-power and MEP amplitude. In neither case, there was an effect of mu-phase on MEP amplitude.

**Conclusion:** The relationship between cortical oscillatory activity and cortical excitability is complex and minor differences in the methodological choices may critically affect sensitivity.

## Introduction

Alpha oscillations (8-12Hz) are a distinct feature of human cortical activity. Initially interpreted to reflect “cortical idling” [1] they are today assumed to have an active role in neural processing: According to the “gating-through-inhibition” hypothesis, alpha oscillations gate information by active inhibition of task irrelevant areas [2, 3]. This hypothesis is supported by studies that demonstrated an increase in alpha power over task-irrelevant cortical areas with simultaneous decrease of alpha power in task-relevant areas [2–5]. Instantaneous fluctuations also actively influence task performance and several studies have shown that performance on perceptual tasks increases, when stimuli are presented during periods of low alpha power [5–7].

While alpha oscillations are most prominent in the occipito-parietal cortex, they are also expressed in pericentral, sensorimotor cortex where they are called mu-rhythm [8–11]. Invasive recordings in monkeys provided evidence that mu-oscillations in the sensorimotor cortex follow the gating-through-inhibition hypothesis: Low sensorimotor mu power predicted an increase in neuronal spiking and increased performance on a tactile perception task [12]. Motor Evoked Potentials (MEPs) induced by single-pulse Transcranial Magnetic Stimulation (spTMS) offer a unique possibility to probe the transsynaptic excitability of fast-conducting corticospinal pathways noninvasively in humans. Several studies used spTMS to examine if the pre-TMS mu-power modulates corticospinal excitability (i.e. the MEP amplitude) using post-hoc grouping of MEPs according to pre-stimulus power. Early studies seem to support the gating-through-inhibition hypothesis and report associations between low mu-power and higher MEP amplitudes [13–15], but these findings could not be replicated by a number of often better-powered studies [16–21] (See [22] for a detailed, tabular report of studies grouping MEPs according to pre-stimulus power). Real-time EEG-triggered stimulation systems makes it possible to assess the relationship between mu-oscillatory activity and MEP amplitude more effectively through online targeting of specific oscillation states [13–15]. In contrast to the studies using post-hoc grouping, several real-time mu-triggered experiments report a positive relationship between mu-power and MEP amplitude, suggesting mu-facilitation rather than a mu-inhibition in the corticospinal system [23–25].

Considering the contradictory evidence from spTMS studies, it is paramount to understand the physiological effects of basic stimulation parameters before the findings of these studies can be used to support or challenge theoretical ideas like the gating-through-inhibition hypothesis. Recently, a real-time EEG-triggered TMS study by Ogata and coworkers found a positive relationship between mu-power and MEP amplitude, but only at higher TMS intensities and when participants had their eyes open. No such relationship emerged at low stimulus intensity or when participants had their eyes closed [24]. This suggests that the relationship between mu-power and corticospinal excitability is not fixed but critically depends on extrinsic (i.e. experimental stimulation and recording set-up) and intrinsic (i.e., brain state) factors.

One important experimental factor is the way how the oscillatory cortical activity of interest is “distilled” from the EEG activity recorded at the scalp level [22, 26, 27]. In this regard, it is noteworthy that all mu-triggered studies, that report a positive linear relationship between mu-power and MEP amplitude used a sensor-level Laplacian montage centered above the M1 [23, 24]. Conversely, post hoc-studies that have shown a negative linear relationship or no relationship at all between mu-power and MEP amplitude have predominantly used source projections or the averaged activity in a cluster of surface electrodes [13, 14, 16, 18, 19, 22].

To re-examine the relation between mu-power and cortico-spinal excitability, as reflected by MEP amplitude, we employed mu-triggered spTMS using an individual source projection to trace pericentral mu-activity. We applied brain-state informed EEG-TMS for real-time power estimation and targeted the highest and lowest 25% of mu-power in each individual. In addition, we also re-referenced our data using a sensor level Laplacian montage centered over M1-HAND and used post-hoc trial sorting, to test whether the EEG montage used to extract power influenced the detected relationship between MEP amplitude and mu power. For both methods, we also defined the pre-stimulus phase and the inter-stimulus interval between two pulses and used these as covariates to explore their ability to predict MEP amplitudes.

## Methods

### Subjects

15 right-handed healthy participants took part in this study (7 female, average age= 24,1 ±2.8 years). The sample size was based on previous studies using state-informed brain stimulation to investigate cortico-spinal excitability [22, 28]. We did not perform any preselection based on individual TMS or EEG characteristics (e.g. size of the single alpha-band peak or resting Motor Threshold (rMT)). Subjects were allowed to enroll in the study if they were right-handed and between 18-40 years of age and did meet the criteria specified in the TMS safety screening questionnaire [29]. All subjects gave informed written consent. The study was approved by the Regional Committee on Health Research and Ethics of the Capitol Region in Denmark and was in accordance with the Helsinki declaration (Protocol H-16017716).

### Experimental setup

Participants were sitting in a relaxed position in a commercially available TMS-chair (MagVenture, Farum, Denmark). Cushioning provided additional arm and neck support and the participant was instructed to keep the hands and arms relaxed and the eyes open throughout the experiment. A real-time EEG-TMS setup was used for the online analysis of the EEG-signal and triggered single TMS pulses when individual mu-power met the predefined criteria of the current stimulation condition (see Figure 1).

**Figure 1:**
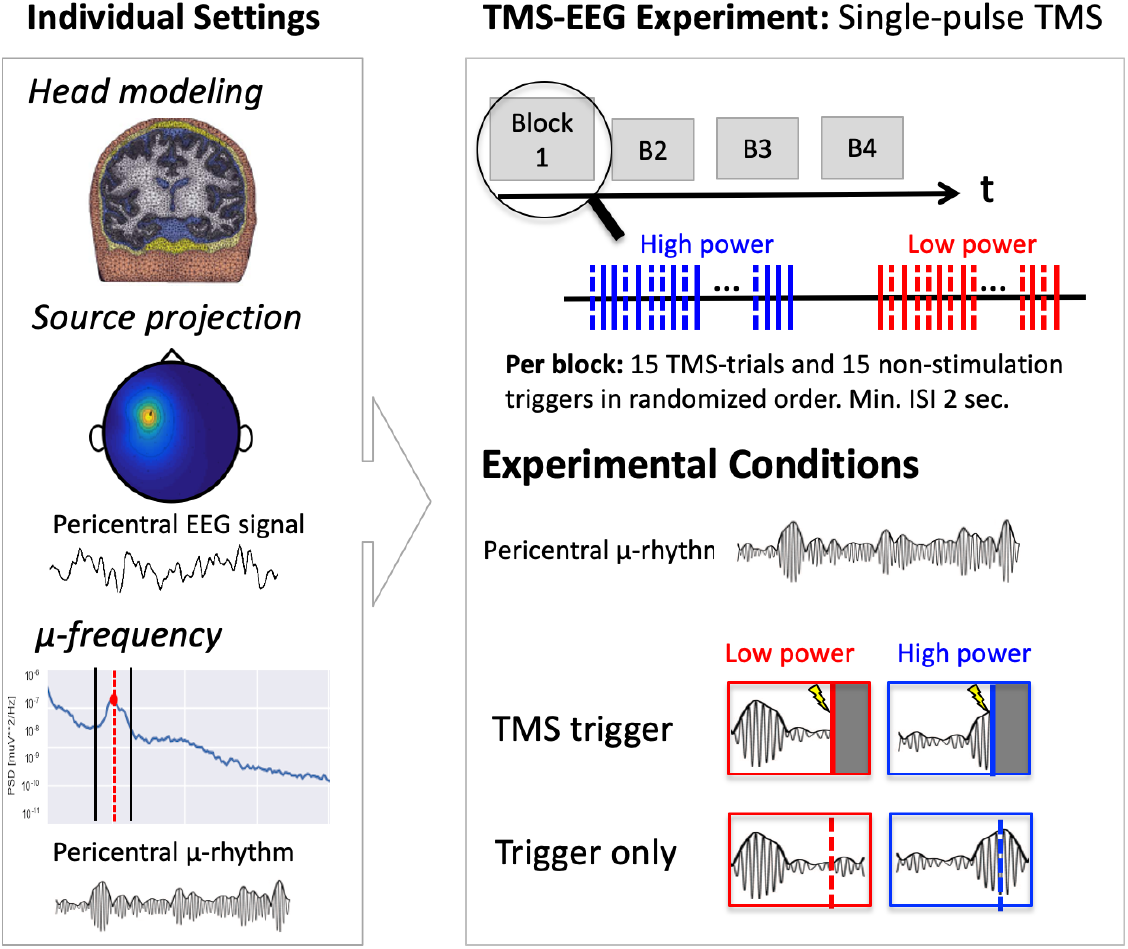
The experimental timeline: The first panel shows how the location and center frequency of the individual mu-rhythm were determined. The second panel illustrates the temporal structure of the TMS experiment and the four conditions.

#### Electrophysiological recordings

Both EEG and EMG were recorded using a NeurOne Tesla system (NeurOne Tesla, Bittium, Oulu, Finland). The amplifiers had a sampling rate of 5kHz and we used a 2.5 kHz antialiasing low-pass filter with 24-bits resolution per channel, across a range of +/−430 mV. To ensure that the latency of the data-delivery for the real-time processing stayed below 5ms, the data was sent directly from the EEG amplifiers Field-Programmable-Gate-Array via a user-datagram protocol (1kHz update rate over a 1Gb/s Ethernet link). The scalp EEG was recorded using a TMS compatible EEG cap (Easycap M10 sintered Ag/AgCl multielectrodes Easy Cap, Woerthsee-Etterschlag, Germany), with equidistant spacing between the 63 surface electrodes. All electrodes were prepared using high-chloride, abrasive electrolyte gel (Easycap, Herrsching, Germany), until the impedance was below 5kΩ. The EMG was recorded using self-adhesive, disposable surface electrodes (NeuroLine, Ambu A/S, Denmark), and the EMG signal was transferred in the same way as the EEG signal described above. The electrodes were applied in a belly-tendon montage on first dorsal interosseous (FDI) muscle of the right hand (the ground was placed on the right wrist), making sure that a clear muscular response was measured and that 50Hz noise was below 20uV.

#### TMS

Single TMS pulses were applied using a figure-of-eight shaped MC-B70 coil connected to a MagPro 100 stimulator (Magventure, Farum, Denmark). The stimulator was set to a monophasic stimulation waveform inducing an A-P current direction in the brain. The motor hotspot (M1-HAND) was defined as the coil position and orientation that resulted in the largest and most reliable MEP amplitude, recorded from the fully relaxed FDI muscle. Stimulation intensity was individually adjusted to elicit a mean MEP amplitude of 1mV, using an in-house threshold hunting algorithm (implemented in Python) based on a previously described adaptive thresholding method [30]. Threshold hunting was initiated at 47% of maximum stimulator output (MSO), and the relative standard deviation of the true threshold was assumed to be 7% [31]. The same procedure was used to estimate the Resting Motor Threshold (RMT) (MEP amplitude >= 50uV). The precise positioning of the TMS coil at the target site (FDI motor hotspot) was continuously monitored using stereotactic neuronavigation (Localite GmbH, Sankt Augustin, Germany).

#### Source projection

A radial source projection was used to extract the pericentral mu-rhythm for each subject. The source projection matrix was calculated using (i) an individual head model based on a structural T1-MRI scan, (ii) the position of the EEG channels, and (iii) the functionally determined FDI motor hotspot. Both (ii) and (iii) were registered on each individual head model using the neuronavigation. The head model was estimated using the boundary-element modelling approach [32] implemented in Fieldtrip (Version date: 2016-01-26). The nasion, as well as right and left tragus were manually marked on the MRI images. Finally, the source space (weighted sum of all EEG electrodes) was estimated using the “dipoli” method implemented in fieldtrip considering a single dipole with radial orientation at the position of the FDI hotspot.

#### Real time estimation of pericentral mu power

During acquisition a ring-buffer containing the latest 500ms of data was updated upon the arrival of each sample. To limit the computational cost of the subsequent processing steps the sampling rate in this buffer was reduced to 1kHz by averaging the 5 samples received in each UDP package. A processing loop running in a separate process with real-time priority performed source projection and a discrete Fourier transform of the 500ms of data [33]. As an indicator of mu-band power we considered the fraction of the sum of the absolute squared coefficients within the defined mu-band and the total signal power, thereby obtaining a mu-band power fraction. The average update time for the processing loop was below 0.5 ms, which ensured that only very occasionally the estimate was not updated for each sample, meaning that the additional latency due to signal processing was negligible.

#### Re-extraction of the pericentral my-rhythm using a Laplacian montage

To test the influence of differences in electrode montages for extracting the mu-power we re-extracted the pericentral mu-rhythm from the raw scalp EEG for each subjects using a sensor level five-channel Laplacian montage centred on the electrode above the M1-HAND area [23]. This enabled the post-hoc determination of pre-stimulus mu-power and mu phase according to a C3-centered Laplacian montage and allowed the direct comparison of source-extracted and surface extracted mu-oscillations.

### Experimental sessions

#### Structural MRI scans

On the day before the experiment each participant was scanned using a structural T1-weighted MRI sequence (T1-w MRI). T1-w scans were done using a magnetization prepared rapid gradient echo (MPRAGE) MRI sequence with 0.85mm isotropic spatial resolution, TR = 6ms, TE = 2.7ms and flip-angle = 8° on a Philips 3T Achieva scanner (Philips, Best, Netherlands). The field of view was 245×245×208mm, such that the scan covered the whole brain. The T1-w scan was required for obtaining the head model for the source projection, as well as for neuronavigation during the state-informed experiment.

#### TMS-EEG Experiment

Each experiment started with a range of pre-measurements to determine the subject specific stimulation criteria. First, the subjects were co-registered with their T1-w scan, then the individual EEG electrode positions were registered to the T1-w scan. The EEG and EMG electrodes were mounted as described above, and the signal quality of both measures were visually monitored throughout the whole experiment, and EEG channels were re-prepared in case the signal quality dropped. The next steps included the determination of the M1-HAND hotspot, the RMT and the stimulation intensity required to evoke MEPs of 1mV (1MV-MEP) amplitude for the right FDI. Both RMT and 1mV-MEP were estimated using the threshold hunting algorithm described in the experimental setup section above. Finally, the source projection matrix was calculated.

To identify the individual mu-rhythm, we recorded 5 minutes of EEG while the subject was resting with open eyes. The individual peak frequency was determined as the peak of the mu-power spectral density (PSD) within 7-13Hz, based on the resting EEG data and the estimated source matrix. The mu-frequency band was defined as ±2Hz around the peak frequency. However, in cases where the lower frequency limit would be below 7Hz, the limit was set to 7Hz, resulting in a narrower frequency band for these subjects. Furthermore, the PSD maps were used to determine the highest (q75) and lowest (q25) quantiles of the mu-power, which later were used for thresholding of the power-informed stimulation.

#### Power-informed stimulation

During the experiment TMS stimulation was triggered by instances of high (q75) and low (q25) power of the endogenous pericentral mu-rhythm, using the thresholds estimated during the pre-measurements. As TMS gives rise to large artefacts in the EEG signal just after the stimulation, a non-stimulation trigger was set in 50% of the trials. This enabled post-hoc evaluation of power estimation performance in the non-stimulated trials, while power-dependent effects of spTMS on corticospinal excitability were evaluated in the 50% of trials with TMS. 60 trials were collected for each of the four conditions (q75-TMS, q75-trigger-only, q25-TMS, q25-trigger-only) resulting in a total of 240 trials, split into 4 blocks. Each block contained 15 trials of each condition, resulting in a total of 60 triggers per block. Stimulation was given in blocks of either high or low power and the order between stimulations and non-stimulations triggers was pseudo-randomized, such that no more than 3 of the same kind could appear in a row.

Between each block, a short break of 1-5 minutes was allowed. For a few subjects, we adjusted the stimulation intensity between blocks, such that the stimulation always elicited an average MEP of around 1mV, however the total adjustment never resulted in a change of more than 4% stimulator output. Average stimulator output across participants was 70% ± 13%. To avoid any systematic interaction between TMS pulses, the minimum inter-trial-interval (ITI) set by the algorithm was 2s. Due to the constraints set in the power-detection algorithm the actual ITI was considerably longer and had a mean of 10.3 s across all individuals. On a few occasions (less than 5% of trials) the ITI either exceeded 60 seconds or was undefined (for the first trial for each block) these trials were removed from further analysis.

### Data Analysis

#### MEP analysis

The peak-to-peak amplitude of the MEP was determined trial-by-trial by an in-house developed python script. Similar to a previously published study [22] all trials with an EMG activity > 50mV during the 100ms prior to stimulation, or with MEP amplitudes more than 2.5 standard derivations away from the mean were excluded from further analysis. On average 6.3% of trials were excluded from further analysis due to these criteria.

#### Phase at Stimulation

The phase at the time point of stimulation was estimated from the recorded data using a continuous Morlet wavelet transform. The transform was done for 51 frequency scales across the mu-band and the phase of the optimal frequency scale 100ms prior to stimulation was projected to the stimulation time using the same algorithmic procedure as described previously [22]. Depending on the estimated phase, all TMS trials were sorted in a post-hoc analysis into four distinct phase bins (0°, 90°, 180°, 270°).

#### Statistics

To test the hypothesis that the cortico-spinal excitability is modulated by mu power at the time of stimulation while simultaneously assessing the relationship between power, phase and interstimulus interval (ISI) on the individual trial basis we performed a mixed-effect analysis that included mu-power fraction (categorical – high, low), the mu-phase (categorical – 0°, 90°, 180°, 270°) and the inter-stimulus-interval between two trials (continuous – ISI) as fixed effects and the participant number as a random effect. Statistical analysis was performed using the statistical software package R (https://www.r-project.org). Mixed effects analysis was performed using the lme4 package (Team RC 2018) and all continuous variables were log-transformed before they were entered in the model. The significance threshold for null hypothesis testing was set to p<0.05.

To directly test evidence for the null hypothesis we additionally used Bayesian analysis of covariance as implemented in Jasp v. 0.11.1 with the MEP as dependent variable, high/low power and phase discretized into four bins as a fixed-factors, subject as a random factor and included the logarithm of the ISI as a covariate.

#### Analysis of the data extracted by Laplacian Surface Montage re-referencing

The analysis performed for source-extracted mu-power were also run for the post-hoc re-referenced Laplacian data. The power in mu-band in the Laplacian montage was only very weakly related to the power in the original source projection meaning that the power fraction could no longer be reasonably divided into two separate bins. Therefore, for the mixed model, we only included the highest and lowest quartile of the Laplacian-power fraction in order to keep the data as similar as possible to the source-triggered analysis, even though this meant bisecting data. However, in the analyses that considered a continuous representation of the power fraction (Bayesian ANCOVA) we utilized all the available data.

## Results

### Online power-triggered EEG-TMS

We verified that the online power triggering worked by assessing the power fraction estimate for the non-stimulated trials in a 500ms window centered at the intended stimulation timepoint. The mean power fraction was 0.15 ± 0.07 in the low-power triggered condition and 0.30 ± 0.12 in the high-power condition, indicating that the pericentral mu-power was successfully targeted by brain-state informed spTMS (Figure 2A). The ISI during the low-power was 13.9 ± 4.2 ms and 10.5 ± 2.0 ms during the high-triggered condition. A paired t-test (unequal variance) testing for differences in the ISI indicated that duration was significantly influenced by the triggering condition (t_14_ = 2.47; p= 0.02) (Figure 2B).

**Figure 2:**
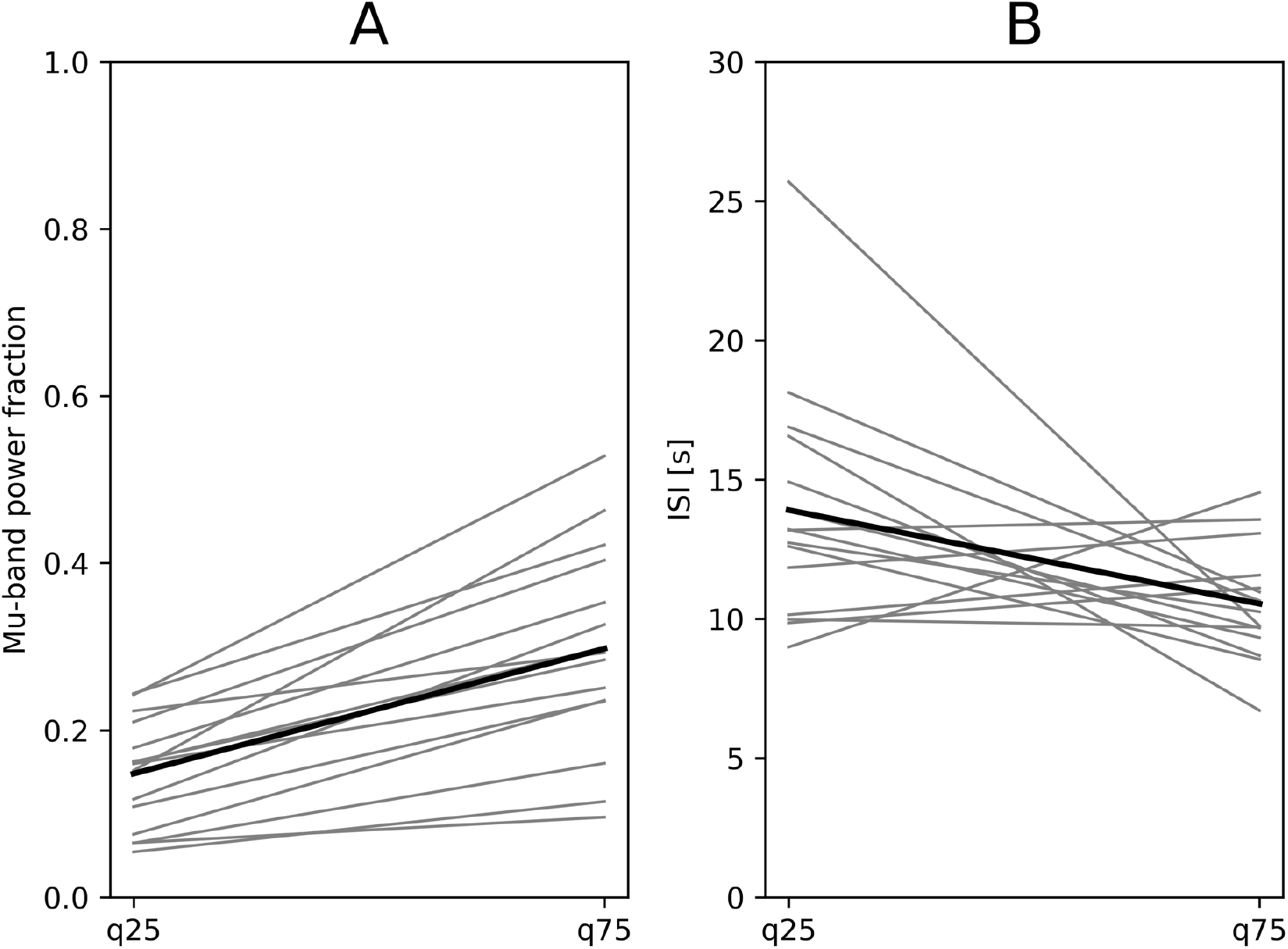
Precision of Targeting: A) shows the power fraction in the mu-band at in the non-stimulated trial, centered at the intended stimulation point in high (q75) and low (q25) power condition. Grey lines represent individuals, the black line the grand average B) shows the average ISI in high (q75) and low (q25) power condition in stimulated trials. Grey lines represent individuals average, the black line the grand average

The accuracy of the phase detection algorithm was evaluated by comparing the projected phase to a centered phase estimate for non-stimulated trials as also described in a previous publications [22]. As expected, this revealed that the phase estimation accuracy was lower for the low power condition (q25) with a mean absolute error of 61°, whereas the mean absolute error in the q75 condition was comparable to our previous work with a value of 47°[22].

### Mu-rhythm extracted by source projection

The average MEP amplitude was 1.16 ± 0.39 mV across both conditions. The mean MEP amplitude in the high-power triggered condition was 1.14 mV compared to 1.17 mV in the low-power triggered condition. The linear mixed-effects model, that treated mu-power and mu-phase and ISI as fixed effects and participant as a random effect showed no significant main effect for power (x^2^(1) = 0.32; p=0.57) (Figure 3A). Also, the main effect of phase was not significant (x^2^(3) =1.54; p=0.20) (Figure 3B), while ISI significantly modulated the MEP amplitude (x^2^(1) = 5.14; p=0.02). None of the interaction terms were significant. The main effect of ISI was caused by an increase of MEP amplitude at when pulses where given at longer intervals (Figure 3C).

**Figure 3:**
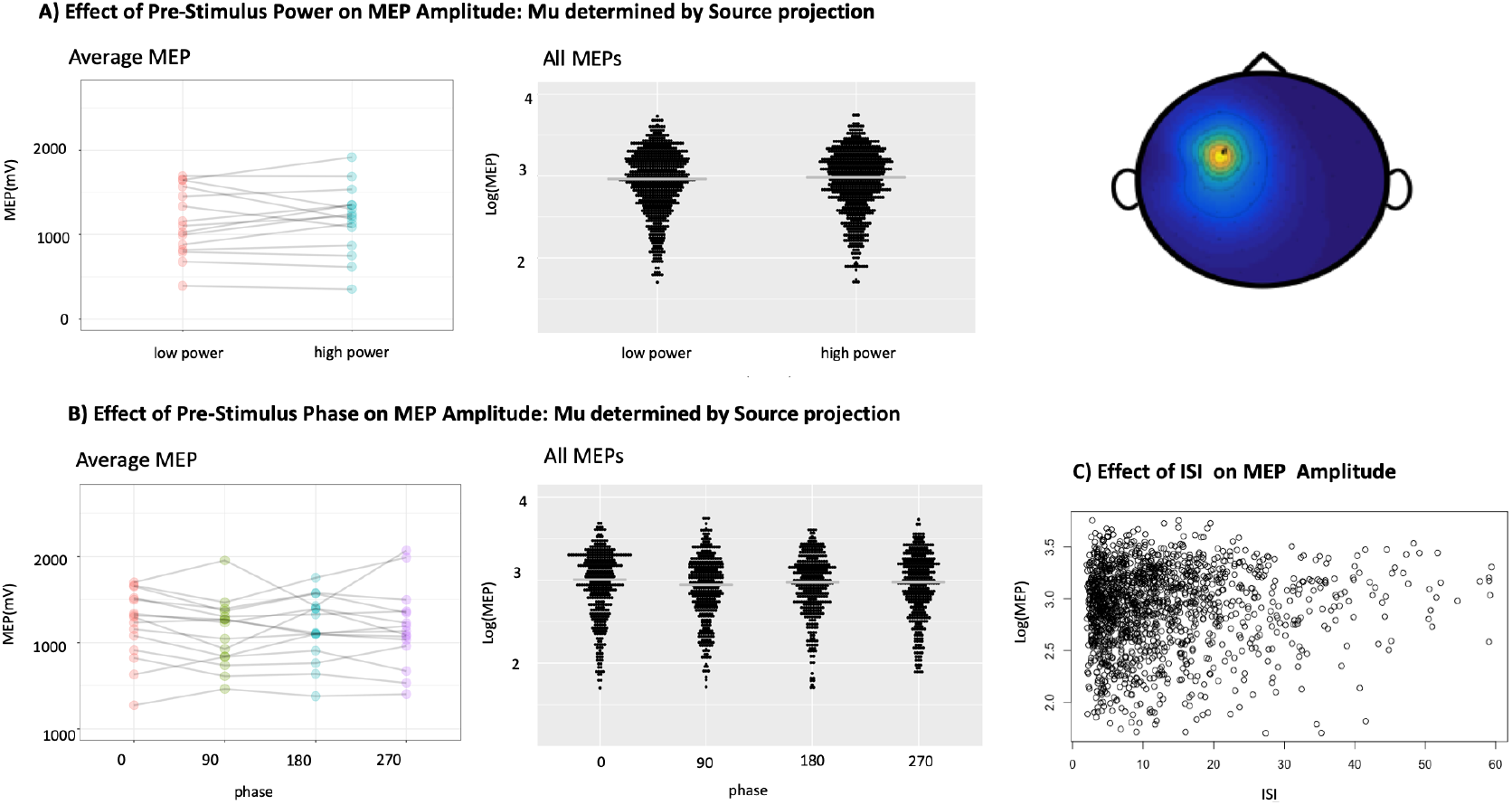
Source Projection: MEP as a function of Power and ISI A) shows the average MEP amplitude at stimulation point in high (q75) and low (q25) power condition. Grey lines represent individuals, the black line the grand average B) shows the relationship between pre-stimulus phase and MEP amplitude and C) shows the relationship between MEP and ISI.

### Bayesian Analysis

Using Bayesian analysis of covariance (ANCOVA) to assess inclusion probabilities for each of the predictors we found no evidence for inclusion of ISI (Bayes Factor (BF): 0.41), strong evidence against inclusion of power fraction (BF: 0.04) and extreme evidence against inclusion of phase (BF=0.006) (Table1).

### Mu-rhythm extracted by Laplacian Surface montage

To test whether the EEG montage used to extract pericentral power influenced the detected relationship between extracted power and cortical excitability, we re-referenced the EEG data using a sensor level Laplacian montage centered on M1-HAND, similar to previous power-triggered experiments [23, 24] and Laplacian mu-oscillations for each TMS pulse were extracted post-hoc. The linear mixed-effects model, that treated Laplacian mu-power fraction and mu-phase and ISI as fixed effects reveal a significant power-effect (x^2^(1) = 5.91; p=0.01) indicating that the MEP amplitude was larger when Laplacian mu-power was high (Figure 4A). For phase, the Laplacian mu-extraction agreed with the initial source-projected data and showed no significant effect of mu-phase (x^2^(3) = 0.56; p=0.64) (Figure 4B). Figure 4 displays the MEP as a function of phase to further illustrate that no effect of phase could be seen. The ISI effect was significant as in the analysis of the source projected data (x^2^(1) = 3.9; p=0.04).

**Figure 4:**
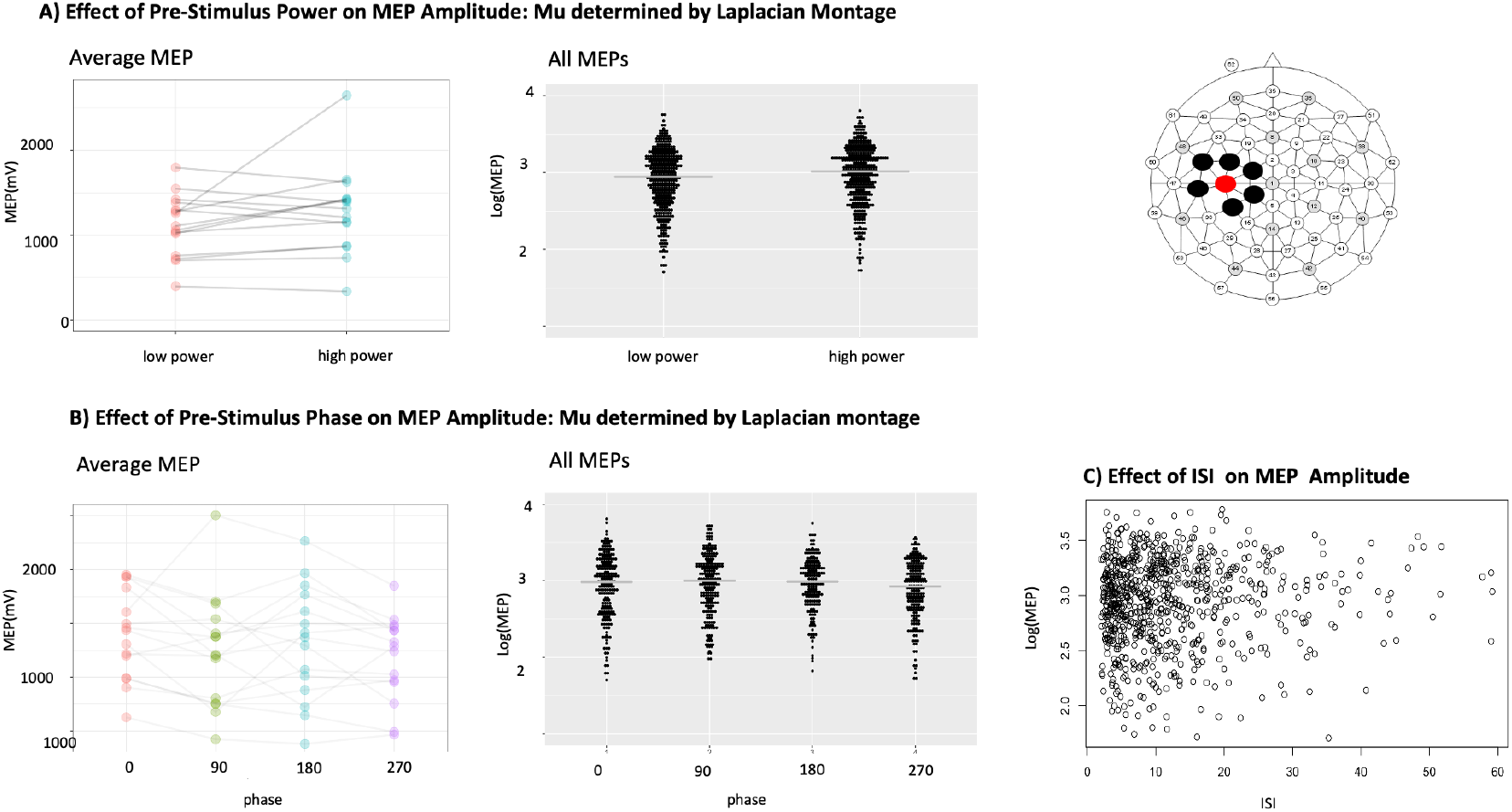
Laplacian montage: MEP as a function of Power and ISI A) shows the average MEP amplitude at stimulation point in high (q75) and low (q25) power condition. Grey lines represent individuals, the black line the grand average B) shows the relationship between pre-stimulus phase and MEP amplitude and C) shows the relationship between MEP and ISI.

### Bayesian Analysis

A Bayesian ANCOVA which considered all the data with the Laplacian montage by including the continuous power fraction as an independent variable. In this analysis the inclusion probabilities showed very strong evidence for inclusion of the power fraction (BF: 75.9) and no evidence for inclusion of ISI (BF: 0.941) and again extreme evidence against the inclusion of phase (BF: 0.002).

### Spatial comparison of radial source and Laplacian montage and pairwise electrode comparisons

The location of the Laplacian montage was more posterior that the radial source at the individual hotspot, as indicated by the mean MNI coordinates (Laplacian montage, electrode 17: [−62, −22, 73]; radial source at hotspot [−55, −13, 85]). The mean displacement from center of gravity (COG) of each montage was similar in both montages (electrode 17: 9.9; hotspot: 9.5 (both MNI mm)). To compare if specific electrodes preferentially contributed to the effect observed in the Laplacian montage, we also calculated mu-power based on the center electrode (electrode above M1-HAND) and each of its neighbors individually (see figure 5B) by using separate Bayesian ANCOVAs with the logarithm of the MEP as dependent variable, subjects as a random factor, the detected phase-bin for the relevant montage as an independent factor and the logarithm of ISI and the relevant power fraction as covariates. We then quantified the effect of power fraction in each of these six electrode pairs at the Bayes factor for inclusion of the continuous power fraction in the model. The results indicated that the significant effect of power was driven by a signal gradient towards the medial/anterior direction in electrode space (difference between electrodes 7 and 17). In all of these models there were extreme evidence against the inclusion of phase (BF<=0.006).

**Figure 5:**
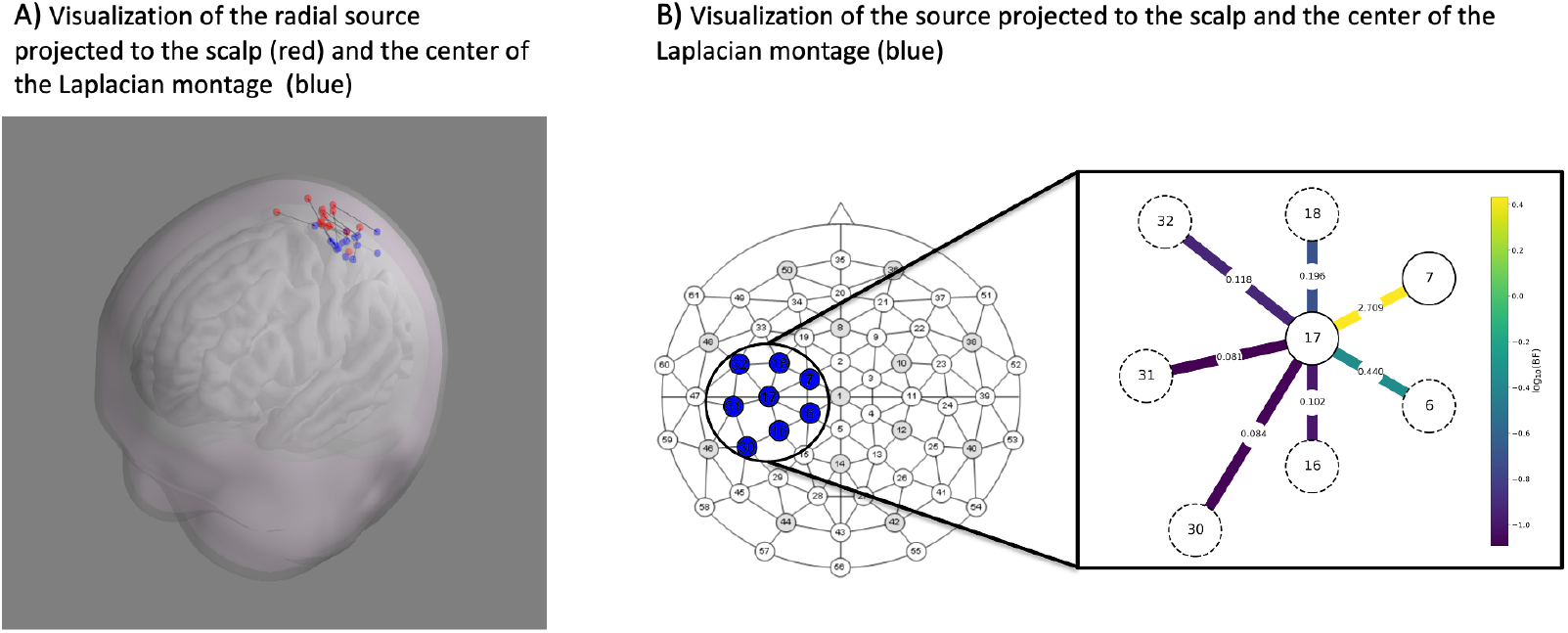
A) shows a visualization of the individual source locations projected to the scalp and the center of the Laplacian montage. Grey lines connect the points for each individual subject. B) Shows the pairwise electrode comparisons in the Laplacian montage checking if specific electrodes preferentially contributed to the observed effect of power. Color coding is according the log-transformed Bayes factor.

**Figure 6:**
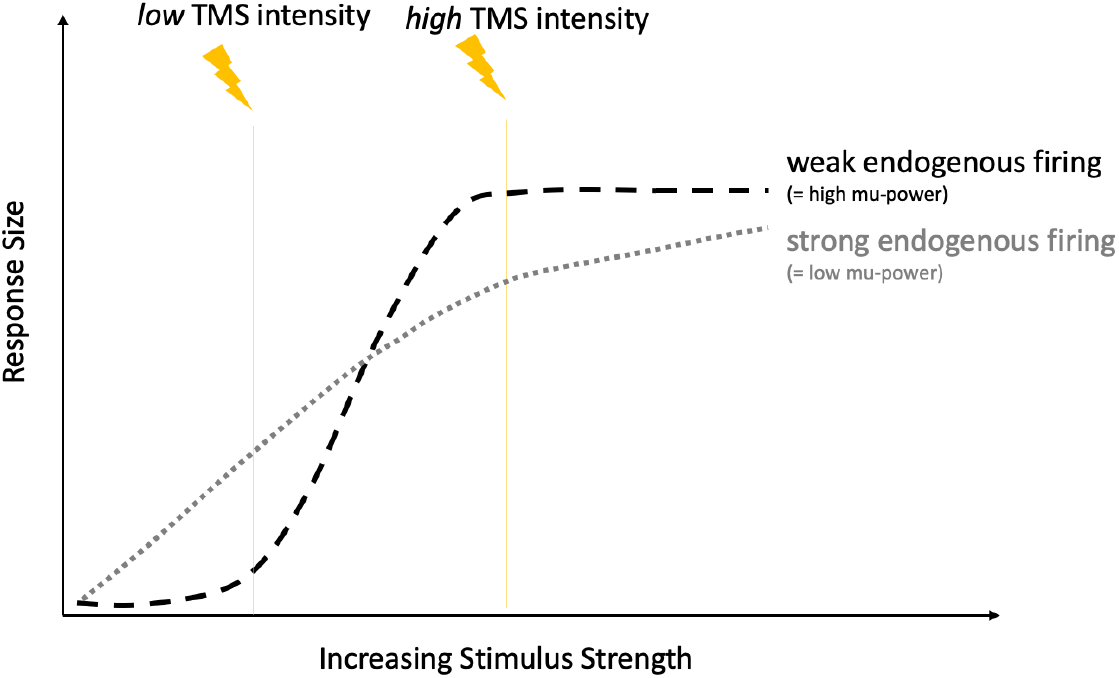
Shows a possible interaction between stimulus strength and background activity. Figure adapted after Matthews. The Effect of Firing on the Excitability of a Model Motor neuron and its Implications for Cortical Stimulation; Journal of Physiology (1999).

## Discussion

Here, we demonstrate that the results of mu-power triggered spTMS are dependent on the electrode montage used to extract the fluctuating expressions of individual pericentral mu-oscillation. A radial source projection, extracting mu-power from the individual motor hotspot for the contralateral FDI muscle, failed to show power-dependent modulation of corticospinal excitability. However, when re-extracting mu-power according to a Laplacian surface montage over M1-HAND, we found a positive relation between pre-stimulation mu-power and MEP amplitude. While the effect of mu-power on MEP amplitude was montage-dependent, both montages concordantly reported no significant effect of mu-phase. Indeed, Bayesian analysis, confirmed that there was strong evidence in favor of the null hypothesis, indicating that mu-phase fluctuations in corticospinal excitability for both montages did not play an important role, thereby replicating the results of our previous work [22]. Our results have important general implications, showing that the way the central EEG signal is extracted from the cortical region of interest will have profound impact on what kind of state-dependent effects of the expressed power or phase in the frequency-band of interest will be revealed by the TMS-EEG approach.

Which property rendered the Laplacian surface montage sensitive to the fraction of pericentral mu-power that has a positive relationship with cortico-spinal excitability? Both, the source projection at the precentral motor hotspot and the Laplacian surface montage were sensitive to radial sources of EEG activity around the central sulcus. The fact that they nonetheless produced different results suggests a complex spatial relationship between pericentral mu-activity and corticospinal excitability and it is interesting to note the spatial differences between the tested montages. The Laplacian montage was more posterior with an average location over the postcentral gyrus (i.e. the somatosensory cortex) while the source projection was on average located over the precentral gyrus (i.e. the primary motor cortex). These results imply that not all sources of pericentral mu-activity in the sensorimotor hand areas modulate cortico-spinal excitability. We cannot exclude that tangentially oriented sources may be relevant as well. Pairwise analysis of the electrodes contributing to the effect seen with the Laplacian montage suggest that a signal gradient towards the medial/anterior direction in electrode space may have contributed to the positive relationship between mu-power and MEP amplitude. Further, more systematic investigations on the impact of different pericentral sources and their spatial location and orientation are required to clarify this question.

Our results fit with the previous body of evidence as studies reporting a positive correlation between pericentral mu-power and MEP amplitude used a surface Laplacian [23, 24] while the studies that used other ways of mu-extraction either reported a negative relationship or no relationship between mu-power and MEP amplitude [14, 18, 22], [13, 16, 19].

How do the present results support the gating-through-inhibition theory of cortical oscillatory activity in the alpha range [2]? The gating-through-inhibition theory would predict a negative relationship between ongoing mu-activity and corticospinal excitability (i.e., MEP amplitude). Apart from a few early reports, all TMS-EEG studies either found no relation or a positive relation which is in apparent contrast to the gating-through-inhibition theory. But maybe the MEP amplitude may not the right measure to test the gating-through-inhibition theory which primarily focuses on cortical processing of sensory inputs and not on generating motor outputs and phase and amplitude of the mu-rhythm may still exert gating-through-inhibition in postcentral sensory cortex. This may or may not have secondary effects on precentral motor cortex which may be opposite in sign, causing corticospinal disinhibition when cortical inhibition in sensory cortex is maximal.

Whatever the relationship between pericentral mu-activity and MEP amplitude may be, any relationship will not be rigidly expressed regardless of the magnitude of cortex stimulation (i.e., the TMS intensity) or the ongoing level of neuronal activity (i.e. the firing rate) [24]. This notion is corroborated by a modelling study using a simple non-compartmental threshold-crossing motoneuron model without dendrites [34]. In that study, Matthews showed that the response of the model neuron to a stimulus depended upon stimulus strength, synaptic membrane noise, and intrinsic tonic firing rate of the neuron (induced by intrinsic background drive). The study revealed a non-linear relationship between background firing rate and the strength of a brief external stimulus: For weak stimuli, the response increased with increasing intrinsic tonic firing rate but for strong stimuli, the response was maximal at a low firing rate and then decreased for higher firing rates. Matthews concluded that “transferring” these findings to corticospinal neurons makes it unlikely that the magnitude of the descending volley elicited by a given cortical stimulus (‘excitability’) will always increase with the initial level of cortical activity. Transferring these observations to TMS-EEG studies, it is unlikely that the MEP amplitude elicited by a TMS pulse will monotonically scale with the level of cortical oscillatory activity (and the associated changes in neuronal firing rates). Instead, strong TMS pulses can be expected to elicit larger MEPs during low levels of intrinsic neural activity (e.g. low firing rate at high mu-power), while weak TMS pulses will elicit larger MEPs during high levels of intrinsic neuronal activity (e.g. high intrinsic firing rate at low mu-power). This flip in the sign of the relationship between mu-power and MEP amplitude is illustrated in Fig. 7. Accordingly, a recent TMS-EEG study demonstrated that high mu-power (e.g. low endogenous activity) was only associated with larger MEP amplitudes during higher TMS intensities [24]. The non-monotonic interaction between intrinsic firing rate (produced by transsynaptic drive) and stimulus response suggested by Matthews [34] also reconcile our present findings (i.e., no relationship or positive relationship between power and MEP depending on montage) with the negative power MEP relationship reported in our recent study [22] in which we only examined the highest power quartile in that work.

Our results have major implications for the use of power-informed TMS-EEG as a tool to better control the cortical “state” at the time of stimulation and hereby, to render the brain response to TMS less variable [35]. Overall, the effect size of the MEP amplitude modulation was modest and sometimes not detectable using conventional average statistics (e.g. ANOVAs)in those studies that reported a relation between pre-stimulus mu-power and MEP amplitude[22–24]. This suggests that the contribution of the pericentral “oscillatory state” to the overall trial-to-trial variability of the MEP is generally low. As long as the interactions between EEG montage, neural background noise, intrinsic firing rate and stimulus intensity are not better understood, the potential of power-triggered TMS to significantly improve the intra-individual variability of single-pulse MEPs remains limited. Additionally, it has to be mentioned that other oscillatory rhythms also have been suggested to modulate cortico-spinal excitability. Especially the sensorimotor beta rhythm but also gamma oscillations and cortico-muscular coherence have also been proposed to modulate cortico-spinal excitability [13, 16, 18–20, 36, 37]. More complex interactions like cross-frequency coupling between cortical oscillation with between difference frequency bands are also possible modulators that have not yet been explored.

Regardless of the mode of extracting pericentral mu-activity, we found no modulatory effects of mu-phase on MEP amplitude. The use of Bayesian statistics also allowed us to move beyond not rejecting the null hypothesis as Bayesian statistics detected strong evidence *against* the contribution of *mu-phase* to fluctuations in corticospinal excitability. Due to a small effect size, effects of mu-power on MEP amplitude may only emerge, if many more (e.g. >100) MEP trials per mu-phase are recorded.

The interval between subsequent TMS pulses is another underexplored aspect of the TMS-EEG approach. In our study, we ensured that TMS was given at very low repetition frequency with a large jitter. This decision was motivated by several TMS studies showing that continuous quasi-repetitive, or jittered application of suprathreshold TMS in the 0.5 – 0.3 Hz range can induce changes in the excitatory-inhibitory balance in M1 and cause change in the inhibitory/faciliatory balance in M1 [38] [39] [40]. Studies also showed that long ISIs (> 0.2 Hz) significantly improve the reliability and lower the variability of intra-individual MEPs [41].

Our experimental procedure resulted in long and highly jittered ISIs, and replicated our previous finding that MEPs tend to be larger when long ISIs are used [22]. This suggests that the preceding pulse may have some conditioning influence on the MEP response produced by the next TMS pulse. These “recency” effects together with the virtue of TMS to shape corticospinal excitability and cortical oscillatory activity, underscore the need that TMS-EEG always need to consider whether or how much the repeated administration of supra-motor threshold TMS pulses “actively” shapes regional cortical excitability and oscillatory activity rather than “passively” probing it.

Taken together, both experimental and theoretical work suggests that the relationship between regional neuronal activity and corticospinal excitability is complex and may interact with a range of different experimental choices such as stimulation intensity, ISI and the number of given stimuli that influence the signal to noise ration and the intrinsic inhibitory/excitatory balance in the stimulated cortex [21, 22, 34, 42, 43]. The relationship between cortical excitability, cortical activity and basic experimental parameters has to be better understood and investigated before brain-state triggered TMs can become a standard technique to reduce inter-trial variability and increase the effectiveness of TMS protocols.

## Acknowledgements

This work was supported by the Novo Nordisk Foundation Interdisciplinary Synergy Program 2014 [“Biophysically adjusted state-informed cortex stimulation (BASICS); NNF14OC0011413]. Hartwig R. Siebner holds a 5-year professorship in precision medicine at the Faculty of Health Sciences and Medicine, University of Copenhagen which is sponsored by the Lundbeck Foundation (Grant Nr. R186-2015-2138).

